# MicroBioAgent: Closed-Loop Planning and Asynchronous Execution for Autonomous Microbiome Analysis

**DOI:** 10.64898/2026.11.17.767465

**Authors:** Zeming Li, Jingjing Wang, Chonghao Wang, Yi Ding, Lu Zhang

**Affiliations:** Department of Computer Science, Hong Kong Baptist University, Hong Kong SAR 999077,China

**Keywords:** microbiome, 16S rRNA, shotgun metagenomics, metagenome assembled genomes, bioinformatics agent

## Abstract

Microbiome data analysis demands customized workflows tailored to specific biological questions, sequencing modalities, and data properties. However, constructing and executing these workflows requires navigating heterogeneous software dependencies, reference databases, and multi-step parameter tuning. Here, we present MicroBioAgent, an autonomous, domain-specific agent framework that translates high-level biological queries into executable, study-specific pipelines across 16S rRNA amplicon, reference- based shotgun, and de novo assembly-based metagenomics. Rather than relying on static pipeline templates, MicroBioAgent employs a closed-loop execution architecture: it dynamically selects and runs bioinformatics tools within user-controlled environments while iteratively inspecting intermediate outputs to adapt downstream analytical decisions. Furthermore, to address the long-running nature of metagenomic computation, we introduce an asynchronous background task-management mechanism that reduces LLM token consumption by over 50% while enabling real-time plan adjustment and automated workflow resumption. We systematically evaluated MicroBioAgent across three benchmarks of increasing complexity: (1) paired amplicon and shotgun profiling of tomato rhizosphere microbiomes, (2) taxonomic and functional profiling coupled with machine-learning classification in a colorectal cancer cohort, and (3) de novo metagenomic assembly, metagenome-assembled genome (MAG) binning, and metabolic pathway reconstruction in a polyethylene degradation study. Across all benchmarks, MicroBioAgent faithfully recapitulated the primary biological findings of the original studies, exhibiting high analytical concordance despite expected tool- and database-dependent variations. In systematic robustness evaluations across 27 independent runs spanning three backbone large language models and three prompt-complexity tiers, MicroBioAgent achieved a 98.8% task completion rate and yielded highly reproducible diversity and differential-abundance estimates. MicroBioAgent establishes a scalable, adaptive paradigm for end-to-end computational microbiome research.

## 1 Introduction

Advances in sequencing have expanded opportunities to investigate microbial communities in human, plant, and environmental systems. Analyses of 16S rRNA gene amplicons and shotgun metagenomes can characterize community composition, reveal functional potential, and recover microbial genomes [1]. Turning these data into biological insights, however, requires researchers to connect sequence processing with taxonomic and functional annotation and downstream statistical analysis. Selecting suitable methods and reference databases, configuring software, writing analysis code, and interpreting the results require both computational skills and biological knowledge. These demands create substantial barriers for researchers without specialist bioinformatics training, limiting their ability to conduct and extend microbiome analyses independently [2, 3].

Existing analysis platforms and workflow tools address parts of this burden. Workflow managers such as Snakemake and Nextflow organize computational dependencies and support reproducible, scalable execution [4, 5]. Once an analysis has been specified, these systems can coordinate its computational steps. Researchers must still determine which analyses address their question, configure the required resources, and interpret the outputs. Adding a new comparison or following up an intermediate finding can also require changes to the workflow or additional code. Reducing this burden therefore requires support for planning and revising analyses as well as executing them.

Large language model agents offer a way to connect these decisions with computational execution through natural-language interaction. AutoBA combined analysis planning, code generation, execution, and automated repair for multi-omics tasks, while Biomni integrated tools and resources for a broad range of biomedical analyses [6, 7]. BioMedAgent, BioMaster, and M3A have further explored automated bioinformatics and human supervision of multi-step analyses [8–10]. For microbiome researchers, the practical need is to carry a study objective through an entire analysis, from processing sequencing reads to comparing communities or examining the functions of individual organisms. Meeting this need requires the agent to select appropriate methods for the input data and connect their outputs to subsequent analyses without requiring users to write commands or assemble the workflow themselves.

MetaClaw is an AI agent designed specifically for microbiome analysis. It connects predefined upstream workflows on FlowHub with customizable downstream analyses through a YAML pipeline registry [11]. This architecture brings upstream processing and downstream interpretation into one agent-guided analysis, while retaining registered workflows and an external platform for upstream metagenomic processing. During execution, MetaClaw manages long-running tasks in the background and monitors their progress through repeated status checks within the active conversation. For researchers who need to combine methods beyond registered workflows or process sequencing data on their own infrastructure, flexible composition and local execution of both upstream and downstream analyses remain important requirements. Flexibility during execution also depends on whether users can revise pending analyses while a computation is running. Keeping the conversation occupied with status checks limits this interaction, motivating an execution mechanism that allows background computation and user feedback to proceed independently.

Here we present MicroBioAgent, an AI-agent framework that translates research objectives into locally executable microbiome analyses. It combines individual analytical skills for 16S amplicon analysis, reference- based metagenomics, and assembly-based metagenomics with downstream statistical analysis and functional annotation, without requiring a predefined end-to-end workflow template. An analysis plan is established before execution and revised in response to intermediate findings or user feedback. Configurable software environments and reference resources support local upstream and downstream execution without uploading raw sequencing reads to an external analysis platform. During long-running computations, the task continues in the background while the conversation remains available for user feedback. The agent can update pending analyses and automatically continue execution after the background task finishes. We evaluated its analytical capabilities through reanalyses of three published datasets comprising 230 sequencing libraries, examining completion of the requested analyses and recovery of reported biological patterns despite differences in methods and databases. To assess robustness, we evaluated task completion and output consistency across three language models and three levels of prompt detail (three replicates per combination; 27 independent sessions). In addition, ablation experiments comparing the system with and without the runtime patch assessed end-to-end task completion, method adherence, and token consumption, as well as whether users could revise pending analyses while a background task was running and whether the agent automatically resumed analysis after the task finished. MicroBioAgent aims to make microbiome analysis more accessible to researchers without specialist bioinformatics training by reducing manual coding, tool integration, and supervision of long-running computation while preserving flexibility in method selection and user control.

## 2 Results

### 2.1 Composing study-specific analyses across microbiome workflows

MicroBioAgent is an AI-agent framework that integrates flexible composition of upstream and downstream microbiome analyses, local execution, and asynchronous human-in-the-loop interaction (figure 1). Given a research question expressed in natural language, it develops an analysis plan and selects microbiome skills to carry it out. These skills provide method-specific guidance that the agent draws on when choosing analytical approaches, preparing commands, and interpreting outputs. As each step completes, the agent examines its results and decides whether the pending plan needs adjustment.

**Figure 1:**
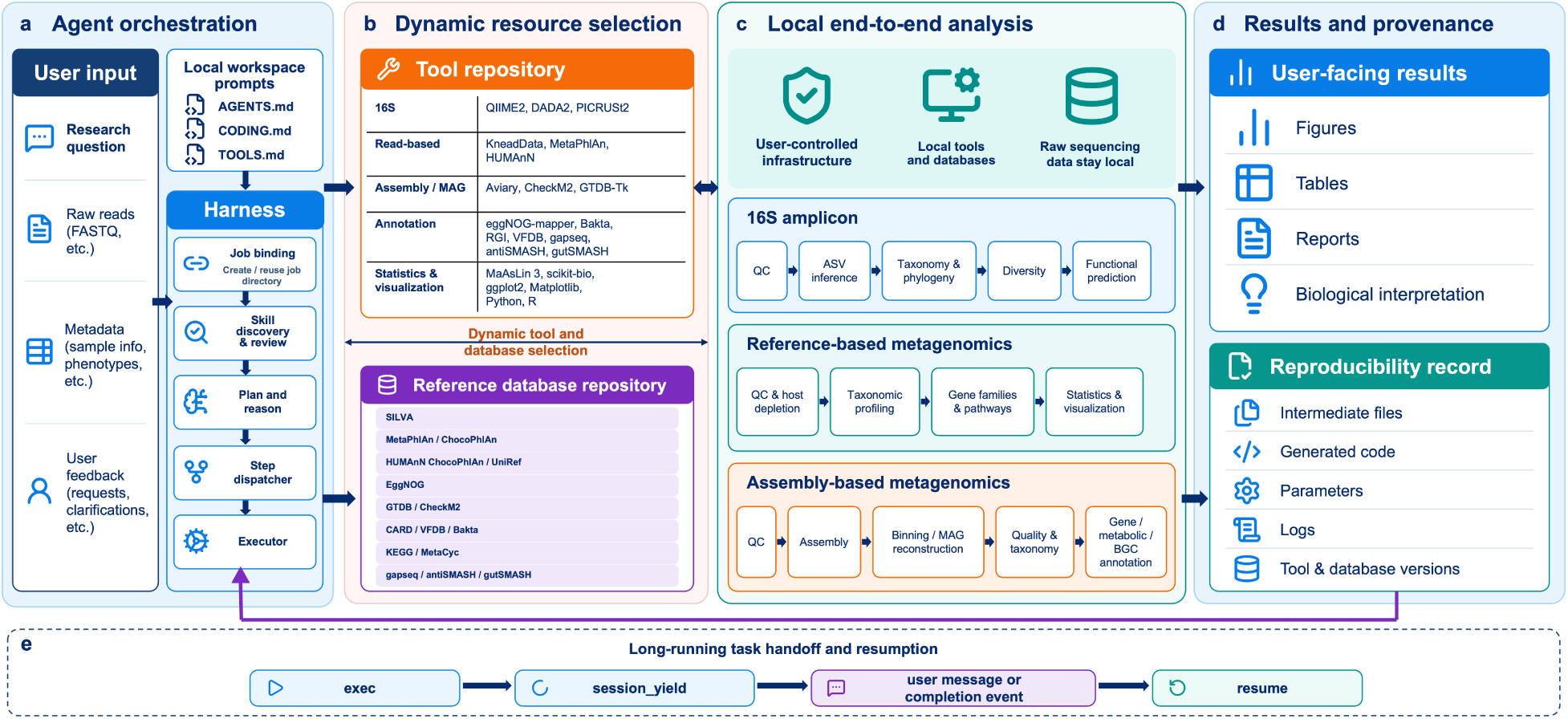
MicroBioAgent architecture and workflow execution. A natural-language research request initiates an analysis workspace, where the agent formulates an analytical plan, selects domain-specific skills, and links local software environments and reference databases across 16S amplicon, reference-based shotgun, and assembly-based metagenomic workflows. Analysis commands, parameters, intermediate tables, and logs are organized within the session directory. During background computation, users can revise pending tasks without interrupting active processes, and task completion automatically triggers output verification and continuation within the same session.

MicroBioAgent executes upstream processing and downstream analyses in a user-controlled local com- putational environment, without requiring raw sequencing data to be uploaded to an external analysis platform. To support these analyses, the system manages analytical tools and reference databases through separate configuration files. The tool configuration specifies available software and its associated execution environments, while the database configuration records reference resources and their local storage paths. Users can modify these configurations to add or replace tools and reference databases according to their research needs.

For upstream analysis, MicroBioAgent supports 16S amplicon processing, reference-based metagenomic profiling, and assembly-based metagenomic reconstruction (table 1). Its amplicon capabilities include sequence quality assessment, denoising, amplicon sequence variant (ASV) inference, taxonomic classification, and phy- logenetic reconstruction. For shotgun data, it supports read quality control and host-read removal, taxonomic and functional profiling, and gene-catalogue construction. Assembly-based capabilities include metage- nomic assembly and binning, quality assessment of metagenome-assembled genomes (MAGs), dereplication, taxonomic assignment, and abundance estimation.

**Table 1:** Overview of analytical capabilities supported by MicroBioAgent across different microbiome task categories.

| Task category | Supported analyses |
|---|---|
| Amplicon characterization | Read import and quality assessment, denoising and ASV inference, taxonomic classification, phylogenetic analysis and functional prediction |
| Shotgun profiling | Read quality control and host depletion, taxonomic profiling, gene-family and pathway abundance profiling, and nonredundant gene catalogue construction |
| Genome-resolved analysis | Metagenomic assembly and binning, MAG quality assessment, dereplication, taxonomic classification, abundance estimation and genome annotation |
| Community analysis | Alpha and beta diversity, ordination, group comparisons, differential-abundance analysis and covariate association testing |
| Functional annotation | Gene and protein annotation, orthology and ontology mapping, functional enrichment, and resistance and virulence gene identification |
| Metabolic reconstruction | Metabolic pathway reconstruction, genome-scale metabolic model reconstruction, and identification of primary metabolic and secondary-metabolite biosynthetic gene clusters |
| Results visualization and reporting | Figures, summary tables and analytical reports |

MicroBioAgent supports a range of downstream analyses, such as comparisons of microbial communities and characterization of the functions of individual organisms. These include comparisons of diversity and taxonomic or functional abundances between groups, covariate association testing, functional enrichment, association network analysis, and supervised classification. For genes and recovered genomes, available analyses include functional and pathway annotation, antimicrobial-resistance and virulence-gene annotation, metabolic reconstruction, and identification of primary metabolic and secondary-metabolite biosynthetic gene clusters. For visualization, MicroBioAgent supports ordination plots of community differences and heatmaps of taxonomic abundances or functional profiles, with customization to highlight the comparisons of interest. During long-running computations, such as MetaPhlAn or HUMAnN profiling, MicroBioAgent keeps the conversation available for user interaction while the analysis runs in the background. Users can provide additional instructions or revise the pending analysis plan without waiting for the running task to finish. This asynchronous human-in-the-loop interaction allows the agent to incorporate feedback during execution and automatically resume the analysis under the updated plan after checking the completed task’s outcome. MicroBioAgent thus supports end-to-end microbiome analysis through flexible workflow composition and local execution, while allowing users to guide ongoing analyses.

### 2.2 Study-specific analyses across plant, human, and environmental microbiomes

We evaluated MicroBioAgent on three published datasets comprising 230 sequencing libraries from plant, human, and environmental microbiomes (Supplementary Table S1). These case studies assessed the system’s ability to complete the requested analyses and recover biological patterns reported in the source studies. Because software versions, reference databases, and some statistical procedures differed (Supplementary Table S2), exact computational reproduction was not the evaluation objective. Instead, we assessed completion of the requested analyses, validity of the resulting outputs, and recovery of prespecified biological signals.

The three cases were selected to test MicroBioAgent on distinct combinations of data types and analytical tasks. The tomato rhizosphere case combined 60 16S amplicon libraries with 36 matched shotgun libraries to examine differences in microbial diversity and taxonomic composition associated with pathogen inoculation and cultivar resistance. The colorectal cancer (CRC) case used 128 faecal shotgun metagenomes from 74 patients and 54 controls to compare species and functional abundances and train classifiers based on gene- family and KEGG Orthology (KO) profiles. The third case used six shotgun metagenomes from soil-derived enrichment cultures grown on low-molecular-weight polyethylene (LMWPE) or alkane substrates to generate MAGs through assembly and binning, dereplicate them, characterize their taxonomic composition, and annotate degradation-associated genes in focal genomes.

#### 2.2.1 A tomato rhizosphere bacterial-wilt resistance study

We applied MicroBioAgent to a published study of tomato rhizosphere microbiomes associated with resistance to bacterial wilt [12]. This case evaluated whether MicroBioAgent could integrate amplicon and shotgun workflows to capture resistance-associated community changes and recover the species markers reported in the original study. The input comprised 60 16S V3–V4 amplicon libraries and 36 matched shotgun libraries from two susceptible cultivars (A57 and HG70), a resistant parent (HG64), and a resistant F1 hybrid (HF12). The amplicon data comprised five sampling conditions with three biological replicates per cultivar: pre-inoculation baseline (T1), and inoculated and uninoculated plants at day 5 (T2 and T2C) and day 10 (T3 and T3C). The matched shotgun libraries covered T1, T2C, and T2. Starting from raw sequencing reads, MicroBioAgent carried out quality control, DADA2 denoising in QIIME 2, and taxonomic assignment against the SILVA database for the amplicon libraries, alongside host filtering and MetaPhlAn3 profiling for the shotgun libraries [13–16].

For the 16S amplicon data, MicroBioAgent examined how pathogen inoculation and cultivar resistance shaped microbial diversity and community composition. Following *Ralstonia* inoculation, Shannon diversity declined in susceptible cultivars but remained higher in resistant cultivars at both day 5 (*P* = 0.002) and day 10 (*P* = 0.004, figure 2a). Community composition also separated clearly by inoculation treatment along the first weighted-UniFrac principal coordinates analysis (PCoA) axis (56.3% of variance; figure 2b)[17]. Separation between resistant and susceptible cultivars was less pronounced along the second axis (12.2%), with partial overlap between groups. Genus-level profiles linked these community shifts to an expansion of *Ralstonia*, which was most pronounced in the susceptible cultivars A57 and HG70 (figure 2c). These patterns were consistent with the inoculation-induced diversity drops and higher post-inoculation diversity in resistant cultivars reported in the original study.

**Figure 2:**
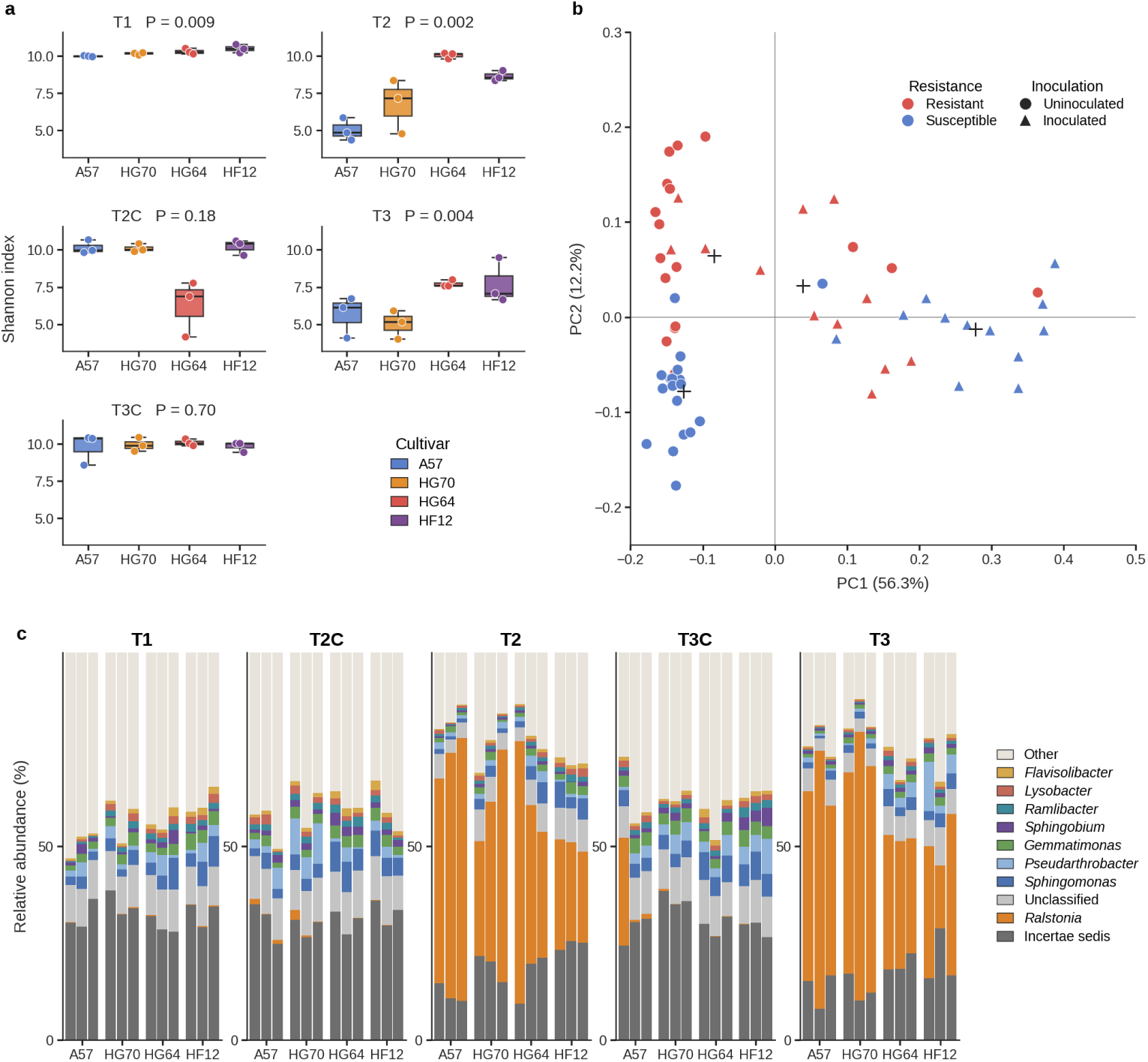
Tomato rhizosphere microbial diversity and community composition. (a) Per-sample Shannon diversity across cultivars, sampling timepoints, and inoculation conditions. (b) Principal coordinates analysis (PCoA) of weighted UniFrac distances, with samples colored by resistance phenotype and point shapes indicating inoculation treatment. (c) Genus-level relative abundance across individual samples. All groups contain *n* = 3 biological replicates (*n* = 60 total 16S samples).

To identify taxa underlying these resistance-associated differences, MicroBioAgent compared each resistant cultivar with the pooled susceptible cultivars using MaAsLin 3 [18]. In the 16S data, between 6 and 15 genera were consistently enriched in both resistant cultivars (HG64 and HF12) across individual sampling conditions (Supplementary Figure S1). To resolve resistance-associated taxa at the species level, MicroBioAgent next analyzed the matched shotgun metagenomic libraries to examine the biomarkers identified in the source study. MetaPhlAn3 profiles showed enrichment of the *Pseudomonas putida* group in both the resistant parent HG64 (*P* = 0.013) and the resistant hybrid HF12 (*P* = 0.009) at T1, but did not detect the other reported marker, *Sphingopyxis unclassified*. Prompted by this difference, the investigator instructed MicroBioAgent to repeat the profiling using MetaPhlAn 2, the version used in the original study [19]. This reanalysis identified both

*P. putida* (*P <* 0.05) and *Sphingopyxis unclassified* (*P <* 0.1, consistent with the significance threshold of the original LEfSe analysis; section 4.2.1) as enriched in the resistant cultivars, recapitulating the published taxonomic biomarker signals (Supplementary Data 1).

#### 2.2.2 A colorectal cancer faecal metagenomic biomarker study

We next evaluated whether MicroBioAgent could integrate taxonomic and functional profiling with downstream biomarker identification, co-occurrence networks, and disease-predictive modeling. The evaluation used a published CRC dataset comprising 128 faecal metagenomes from the Hong Kong C1 cohort, including 74 patients with CRC and 54 controls [20]. Starting from sequencing data, MicroBioAgent carried out species-level profiling with MetaPhlAn3 and quantified UniRef90 gene-family and KO abundances using HUMAnN3 (figure 3)[16]. These profiles served as inputs for downstream community comparisons, biomarker discovery, and cross-validated classification.

**Figure 3:**
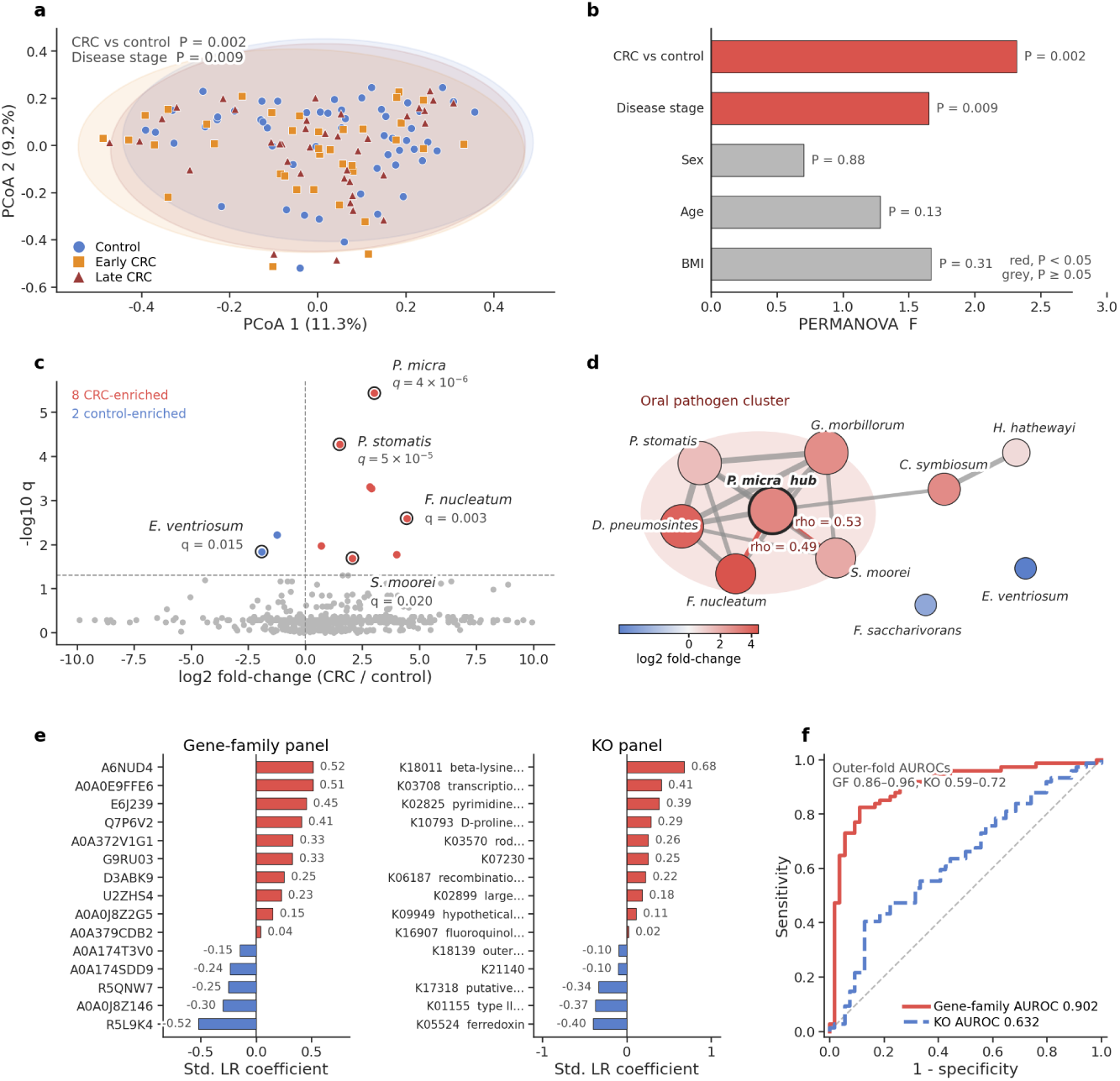
Metagenomic profiling, biomarker discovery, and predictive modeling in the CRC cohort. (a) PCoA of species-level Bray–Curtis dissimilarities across controls, early-stage CRC, and late-stage CRC cases. (b) PERMANOVA test statistics (*F* values) for disease status, stage, and clinical covariates (sex, age, and BMI). (c) Volcano plot of differentially abundant species between CRC cases and controls (Benjamini– Hochberg adjusted *q <* 0.05). (d) Spearman co-occurrence network of the ten differentially abundant species (*|ρ| ≥* 0.4, *q <* 0.05), highlighting the *Parvimonas micra* hub. (e) Standardized logistic regression coefficients for the top 15 gene-family and top 15 KO features selected by mRMR within training folds. (f) ROC curves from nested fivefold cross-validation comparing gene-family and KO classifiers.

PCoA of Bray–Curtis dissimilarities derived from species relative abundances showed no distinct separation among controls, early-stage CRC cases (*n* = 36), and late-stage CRC cases (*n* = 38). A permutational multivariate analysis of variance (PERMANOVA) nevertheless detected significant differences in community composition associated with CRC status (*F* = 2.32, *P* = 0.002) and disease stage (*F* = 1.65, *P* = 0.009)[21]. Potential confounding factors, including sex, age, and body mass index, showed no significant associations with community structure.

Differential abundance analysis between cases and controls identified eight CRC-enriched and two control- enriched species (Benjamini–Hochberg adjusted *q <* 0.05). Four key CRC-enriched species—*Parvimonas micra*, *Peptostreptococcus stomatis*, *Fusobacterium nucleatum*, and *Solobacterium moorei*—and one control-enriched species, *Eubacterium ventriosum*, showed the same direction of association reported in the original study. To determine whether these disease-associated taxa co-occurred across samples, MicroBioAgent constructed a Spearman correlation network from the ten differentially abundant species. The resulting network comprised 16 positive correlations (*|ρ| ≥* 0.4, *q <* 0.05), with *P. micra* acting as a central hub connected to six other species. In particular, its positive correlations with *F. nucleatum* (*ρ* = 0.49) and *S. moorei* (*ρ* = 0.53) recapitulated key co-occurrence patterns highlighted in the source study [20].

Beyond taxonomic markers, MicroBioAgent evaluated whether functional profiles could accurately distin- guish CRC cases from controls. Using minimum redundancy maximum relevance (mRMR) feature selection within a nested fivefold cross-validation framework, the agent trained logistic regression classifiers on the top 15 gene-family or top 15 KO features per fold [22]. The gene-family classifier achieved a pooled out-of-fold area under the receiver operating characteristic curve (AUROC) of 0.902, substantially outperforming the KO classifier (AUROC 0.632). At decision thresholds chosen by Youden’s index, the gene-family model achieved a sensitivity of 0.82 and a specificity of 0.89, compared with 0.41 and 0.87 for the KO model, confirming that fine-grained gene-family abundances provided superior diagnostic discrimination (Supplementary Data 1).

#### 2.2.3 A polyethylene-enrichment genome-resolved metagenomic study

We next evaluated whether MicroBioAgent could reconstruct MAGs from enrichment cultures, quantify their community abundances, and annotate metabolic pathways for alkane terminal oxidation, 2-ketone or oxidized-derivative metabolism, and *β*-oxidation. The dataset comprised six Oxford Nanopore long- read metagenomes from two soil inocula (S1 and S2), each enriched on LMWPE, triacontane (C30), or tetracontane (C40) [23]. Starting from long-read data, assembly and binning with Aviary yielded 372 bins, of which 153 met CheckM2 quality thresholds for completeness and contamination (100 high-quality and 53 medium-quality MAGs)[24–26]. Dereplication at 95% average nucleotide identity (ANI) retained 75 non-redundant representative genomes (50 high quality and 25 medium quality; figure 4a). MicroBioAgent assigned taxonomic classifications using GTDB-Tk (release R232) and estimated MAG relative abundances across the six cultures via CoverM read mapping [27, 28].

**Figure 4:**
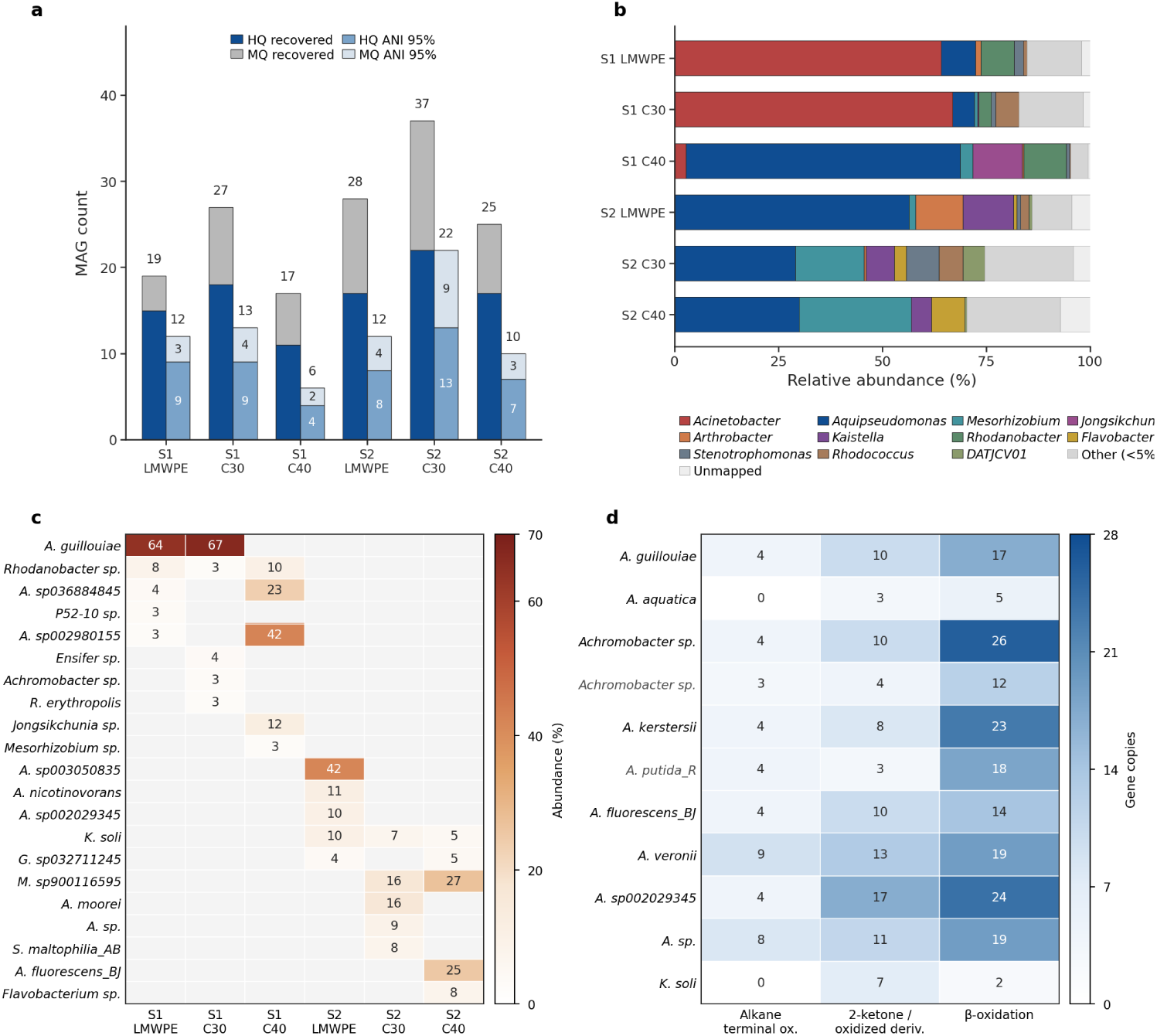
Genome-resolved metagenomic analysis of polyethylene-enrichment cultures. (a) Counts of high- quality (HQ) and medium-quality (MQ) MAGs before and after 95% ANI dereplication. (b) Genus-level relative abundance of MAGs across the six enrichment cultures. (c) Relative abundances of species ranked in the top five of any of the six enrichment cultures. (d) Gene copy numbers annotated by eggNOG-mapper in 95% ANI representative MAGs of focal taxa for alkane terminal oxidation, 2-ketone or oxidized-derivative metabolism, and *β*-oxidation; taxa in gray font denote medium-quality MAGs.

MAG abundance profiles revealed distinct dominant taxa across the enrichment conditions (figure 4b,c). In inoculum S1, *Acinetobacter guillouiae* predominated in LMWPE (64%) and C30 (67%), whereas *Aquipseu- domonas* dominated C40 (66% genus-level abundance). The S2 cultures displayed distinct profiles: *Aquipseu- domonas* sp003050835 was most abundant in S2 LMWPE (42%), *Mesorhizobium_M* sp900116595 and *Aquipseudomonas moorei* led in S2 C30 (16% each), and *Mesorhizobium_M* sp900116595 and *Aquipseu- domonas fluorescens_BJ* predominated in S2 C40 (27% and 25%, respectively). MicroBioAgent successfully recovered all ten focal taxonomic groups highlighted in the source study (Supplementary Data 1). Taxa designated as *Pseudomonas* in the original publication were classified here as *Aquipseudomonas*, consistent with updated GTDB taxonomic boundaries.

To characterize the metabolic capabilities of these recovered organisms, MicroBioAgent annotated the 95% ANI representative genomes corresponding to the focal taxa with eggNOG-mapper [29, 30]. Because *Achromobacter* sp. formed two distinct ANI clusters, both representatives were included, yielding 11 analyzed MAGs (figure 4d). Genes for alkane terminal oxidation were identified in nine of these MAGs, while genes for 2-ketone or oxidized-derivative metabolism and the *β*-oxidation module were detected in all 11 genomes (Supplementary Data 1).

### 2.3 Robustness across models and prompt granularity

To assess whether MicroBioAgent maintains reliable execution and consistent analytical outputs across lan- guage models and levels of user guidance, we crossed three models—deepseek-v4-pro, deepseek-v4-flash and glm5.3-flash—with detailed, medium and simple prompts on the preprocessed Case 1 dataset (3 *×* 3 *×* 3 = 27 independent sessions). Across six core analyses (16S alpha diversity, 16S beta diver- sity, 16S genus-level taxonomic profiling, 16S differential abundance, shotgun species-level taxonomic profiling and shotgun differential abundance), MicroBioAgent completed 160 of 162 evaluations (98.8%; Figure 5a). deepseek-v4-pro and deepseek-v4-flash completed all six analyses in three of three sessions under every prompt level, and glm5.3-flash completed all 16S analyses (9/9 sessions) and 8 of 9 shotgun sessions, with one session under the detailed prompt not analysing the shotgun libraries.

**Figure 5:**
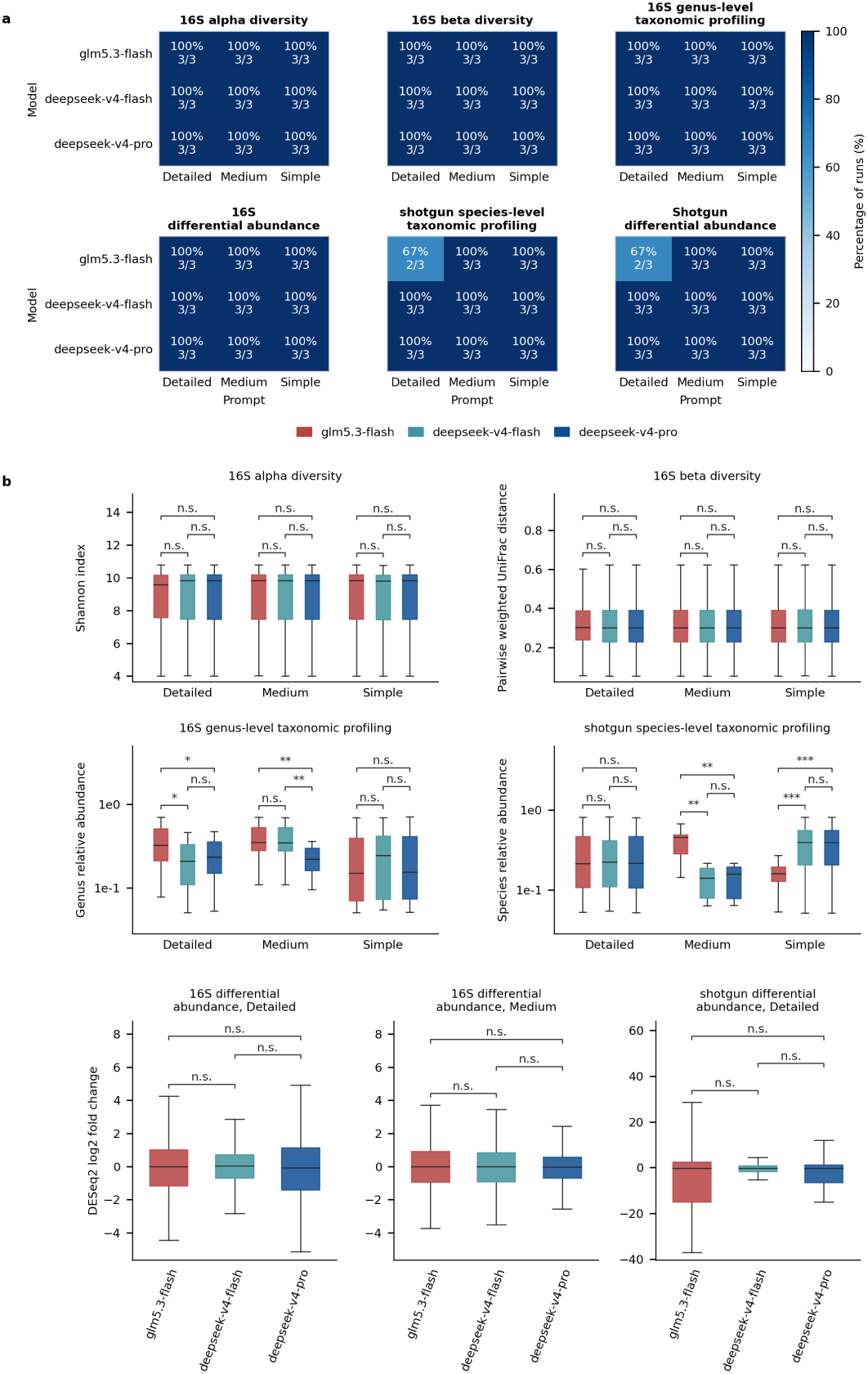
Overall task completion and differences in analytical outputs across models within each prompt level. (a) Completion of six analyses by model and prompt level; values represent completed sessions out of three. (b) Comparison of 16S Shannon diversity indices, pairwise weighted UniFrac distances, and relative abundances (16S genus level and shotgun species level) among models under each prompt level, along with DESeq2 log_2_ fold changes in taxon abundance for 16S analyses (detailed and medium prompts) and shotgun analyses (detailed prompt). Brackets indicate two-sided Mann–Whitney *U* tests with Holm correction across the three model pairs in each plot. n.s., adjusted *P ≥* 0.05; \**P <* 0.05; \*\**P <* 0.01; \*\*\**P <* 0.001. Values are derived from repeated analyses of the same Case 1 dataset and do not represent independent biological replicates.

Community diversity and differential-abundance estimates exhibited broadly concordant distributions across models and prompt levels (Figure 5b and Supplementary Figure S2). Per-sample 16S Shannon indices (median 9.59–9.83) and pairwise weighted UniFrac distances (median 0.301–0.303) did not differ between any pair of models under any prompt level or across prompt levels within any model (all Holm-adjusted *P* = 1, two-sided Mann–Whitney *U* test). For differential abundance, because none of the prompts specified a statistical test, individual sessions chose among DESeq2, MaAsLin 3, Wilcoxon rank-sum tests, Mann–Whitney *U* tests and LEfSe (Supplementary Tables S4 and S5)[31, 32]. Because different tests define effect sizes on different scales, we compared log_2_ fold-change distributions across sessions that used the same test. Across models within each prompt level, DESeq2 was used by all three models for 16S analyses under the detailed and medium prompts and for shotgun analyses under the detailed prompt, and its log_2_ fold-change distributions did not differ between any pair of models (all Holm-adjusted *P ≥* 0.059; under the medium prompt in shotgun analyses, no single test was used by all three models). Between the detailed and medium prompts within each model, log_2_ fold-change distributions also did not differ for any model in either 16S analyses using DESeq2 (deepseek-v4-pro, *P* = 0.54; deepseek-v4-flash, *P* = 0.50; glm5.3-flash, *P* = 0.13) or shotgun analyses using the test shared within each model (deepseek-v4-pro, Wilcoxon rank-sum test, *P* = 0.58; deepseek-v4-flash, Mann–Whitney *U* test, *P* = 0.24; glm5.3-flash, DESeq2, *P* = 0.064).

For taxonomic profiling, relative abundances were more sensitive to model and prompt choices (Figure 5b and Supplementary Figure S2). Across models, relative-abundance distributions did not differ in 16S analyses under the simple prompt or in shotgun analyses under the detailed prompt (all Holm-adjusted *P* = 1), but differed in the remaining prompt settings (Holm-adjusted *P ≤* 0.05). Across prompt levels within each model, distributions were consistent across all three prompts for glm5.3-flash in 16S analyses and for both deepseek-v4-pro and deepseek-v4-flash in shotgun analyses (all Holm-adjusted *P* = 1), whereas differences occurred in the remaining comparisons (Holm-adjusted *P ≤* 0.025). Inspecting the session logs showed three causes of these differences: (1) some sessions applied a V4 rather than a V3–V4 classifier during 16S taxonomic classification; (2) an occasional script misaligned feature identifiers when merging sample profiles into a taxonomic abundance table; and (3) without prompt constraints on MetaPhlAn parameters, sessions differed in whether –unknown_estimation was enabled, changing whether species relative abundances were normalized to total reads or only classified reads. Overall, MicroBioAgent maintained high execution completion and broadly reproducible diversity and differential-abundance profiles across models and prompt settings, while variations in taxonomic relative abundances highlighted the sensitivity of downstream outputs to autonomous tool and parameter selections.

### 2.4 Evaluating asynchronous execution and guidance

#### 2.4.1 End-to-end execution and token consumption

To evaluate how background-execution coordination affects end-to-end analysis, we compared the complete system (full) against an ablated configuration without the runtime patch (no-patch; Supplementary Table S6 and section 4.1.3). Both conditions used deepseek-v4-pro, the detailed Case 1 prompt, and the preprocessed Case 1 dataset across three independent sessions, with no user messages after initial plan confirmation.

Across the three sessions, the complete system produced all 27 requested outputs (9 of 9 per session) and followed all 24 prespecified methodological requirements (8 of 8 per session). The no-patch configuration followed all 24 methodological requirements and produced 26 of 27 requested outputs, with one session omitting archived analysis scripts. Removing asynchronous execution coordination therefore did not prevent the model from completing the prespecified downstream workflow when the prompt explicitly named the required methods, but it substantially altered how long-running commands were managed.

In the complete system, once a long-running command was moved to the background, the agent paused its conversation turn (via the sessions_yield tool, mean 11.7 calls per session, range 5–17) to await automatic notification upon command completion, and rarely queried whether the background process had finished (via the process tool, mean 7.3 calls, range 0–21, with two sessions making at most one call). In no-patch, by contrast, the agent never paused to await completion notifications (0 sessions_yield calls across all three sessions) and instead repeatedly queried the status of running commands with process (mean 155.0 calls per session, range 102–227). Because each status check required a new model call that re-read the growing conversation history, total model calls increased from a mean of 133.3 (range 124–150) in full to 286.3 (range 211–403) in no-patch, and mean total token consumption rose by 141.5% (2.4-fold), from 14.53 *×* 10^6^ (range 12.92–15.84 *×* 10^6^) to 35.08 *×* 10^6^ (range 23.77–49.32 *×* 10^6^) tokens per session (Supplementary Table S6).

#### 2.4.2 Asynchronous human guidance during active computation

We next tested whether the user could revise the pending analysis plan while a minutes-scale Case 1 QIIME 2 taxonomy-classification command was running in the background. Once the classification process was confirmed active, the user sent an instruction to keep the running classification command and update the pending plan to use DESeq2 for the subsequent differential-abundance analysis and generate a heatmap of significant genera.

In the complete system (full), after starting the classification task, the agent paused its conversation turn via sessions_yield and updated the pending plan in response to the user’s instruction while the task was still running (3*/*3 sessions; Supplementary Table S7). Once the classification task finished, the session was automatically reactivated to continue the analysis without additional user input or repeated process checks (3*/*3 sessions). In no-patch, by contrast, because the agent did not pause its turn via sessions_yield and instead kept the conversation occupied with repeated process calls to check whether the task had completed, neither mid-run plan revision nor automatic session reactivation upon command completion was achieved in any of the three sessions (0*/*3).

## 3 Discussion

MicroBioAgent combines modular domain skills, local execution environments, and asynchronous human- in-the-loop interaction to translate biological research questions into executable microbiome analyses. This framework is supported by three complementary lines of evidence: (1) three case studies across plant, human, and environmental microbiomes demonstrating flexible composition across amplicon, shotgun metagenomic, and genome-resolved workflows; (2) 27 independent sessions across three language models and three prompt granularities establishing execution robustness and quantitative output consistency; and (3) ablation experi- ments demonstrating that the runtime patch substantially reduces token overhead while enabling mid-run plan revision and automatic continuation. By coordinating established bioinformatics tools within an inspectable and interactive workflow, MicroBioAgent allows researchers to transparently steer complex analyses without manual coding or fixed pipeline constraints.

The three case studies demonstrate how MicroBioAgent adapts to distinct research questions, spanning cultivar-associated marker discovery in paired amplicon and shotgun libraries, disease-predictive modeling from functional profiles, and genome-resolved metabolic reconstruction. At the same time, these reanalyses show that biological findings remain sensitive to statistical thresholds, tool configurations, and reference database versions. In the tomato rhizosphere study, initial profiling with MetaPhlAn3 detected only one of the two reported species markers, whereas repeating the analysis with MetaPhlAn2, the version used in the original study, recovered both. By preserving commands, parameters, and intermediate tables within the session context, MicroBioAgent makes these analytical choices explicit, allowing researchers to inspect the results and adjust the workflow when needed.

Several biomedical and multi-omics AI agents have been recently developed to automate computational workflows, including BIA [33], BioChatter [34], AutoBA [6], Biomni [7], BioAgents [35], M3A [10], BioMedA- gent [8], BioMaster [9], and MetaClaw [11]. To clarify the distinct capabilities and scope of MicroBioAgent within this evolving landscape, we conducted a systematic functional comparison with these systems (Supple- mentary Table S3). General-purpose agents such as Biomni combine tool retrieval with code-based execution across diverse domains [7], but lack the domain-specific skill guidance and preconfigured reference databases required for end-to-end microbiome analyses beginning directly from raw sequencing reads. Within the microbiome domain, MetaClaw spans upstream and downstream analyses by coupling predefined FlowHub workflows with customizable downstream skills [11]. In contrast, MicroBioAgent unifies both upstream processing and downstream analysis into modular skills that run entirely on the user’s local infrastructure. This on-premises architecture keeps raw sequencing data secure, eliminates dependencies on external comput- ing platforms, and allows method choices to be inspected and adjusted at any stage. Such an interactive approach complements workflow managers such as Nextflow or Snakemake rather than replacing them. While traditional workflow engines excel at large-scale, deterministic execution of static pipelines, MicroBioAgent is tailored for exploratory and iterative analyses where plans could be guided and refined in response to intermediate results.

The cross-model and cross-prompt benchmark illustrates the effectiveness of MicroBioAgent in automating complex microbiome workflows. By generating custom scripts, switching software environments, and coordinating input–output handoffs across tools, the system achieved high overall task completion (98.8%) and consistent diversity and differential-abundance distributions across models and prompt granularities on the preprocessed Case 1 dataset. Nevertheless, occasional inaccuracies in model reasoning also oc- curred, including misidentifying the V3–V4 amplicon region as V4, altering MetaPhlAn normalization flags (–unknown_estimation), and merging feature tables by row position rather than matching identifiers. As foundation model capabilities continue to advance, such procedural inconsistencies are expected to diminish. More fundamentally, these findings highlight the practical value of MicroBioAgent’s transparent architecture. By automating end-to-end computational execution while keeping intermediate commands, parameters, and outputs fully inspectable, the system enables researchers to readily verify analytical decisions and refine workflows without constructing them manually.

The ablation experiments evaluate the operational contribution of the runtime patch during long-running microbiome computations (Supplementary Tables S6 and S7). In conventional agent setups without the patch, monitoring an active process occupies the conversational turn with repeated status checks, which increased mean model calls from 133.3 to 286.3 and inflated total token consumption by 2.4-fold (from 14.53 *×* 10^6^ to 35.08 *×* 10^6^ tokens per session) due to continuous re-reading of the expanding conversation history. More importantly, keeping the session busy with status polling blocks user interaction, preventing researchers from providing timely guidance while computations are underway. By pausing the conversation turn and relying on completion events to reactivate the session, the runtime patch eliminates status-polling overhead while enabling reliable mid-run plan revision and automatic continuation. This capability is especially valuable for multi-step microbiome analyses, where heavy operations such as sequence denoising, taxonomic classification, and metagenomic assembly often run for hours while researchers review intermediate findings and refine downstream hypotheses.

While the case studies and benchmarks show that MicroBioAgent can support interactive microbiome analyses, several practical limits remain. First, our robustness and ablation experiments used the preprocessed Case 1 dataset, and mid-run guidance was tested on a minutes-scale 16S taxonomic classification step across three replicates per condition. Testing the system on multi-hour assembly pipelines, broader command failures, and more detailed plan changes remains an important next step. Second, automated execution checks do not guarantee that biological findings are correctly interpreted, so researchers still need to inspect intermediate tables and confirm overall analysis plans. Third, running analyses locally protects sensitive sequencing data, but users must still install and maintain the required tools, Conda environments, and reference databases on their own machines. Future work should include direct benchmark comparisons with a general coding agent to isolate the practical contribution of preconfigured domain skills and runtime setups. In addition, formal user studies will be valuable to assess whether this conversational framework measurably reduces operational time and cognitive burden for life scientists. Finally, while this study focused on amplicon and metagenomic data, the core architecture of combining modular skills, local environments, and background execution can be extended to other areas, such as transcriptomics or metabolomics, by adding new skill guides and software environments. Overall, MicroBioAgent is not intended to replace human expertise, but to provide a transparent assistant that handles repetitive bioinformatics execution while leaving researchers in full control of analytical decisions.

## 4 Methods

### 4.1 Details on implementing MicroBioAgent

MicroBioAgent was implemented on OpenClaw to plan and execute microbiome analyses from natural- language requests (figure 1). Users communicate with the system through a web interface, a command-line client, or supported messaging applications such as WhatsApp, Slack and Telegram. These interfaces send user requests to the OpenClaw gateway, which maintains the conversation and passes the messages to the language model. When the model requests a tool operation, the gateway invokes the corresponding tool. For computational analyses, bioinformatics commands and statistical scripts run on the user’s workstation or server, using software installed directly on the host or within a docker container. Execution status and captured output return to the conversation, allowing the model to determine subsequent actions.

The behaviour of MicroBioAgent is guided by Markdown instruction files in its agent workspace. AGENTS.md governs task planning and progress tracking, while TOOLS.md describes access to software and reference resources. CODING.md guides code execution and output inspection. Method-specific skills provide analytical guidance, and separate tool and database configurations identify the resources used to execute each step. Supporting user interaction during long-running steps also requires coordination between command execution and the conversation. We therefore introduced a patch to the OpenClaw runtime that modifies tool-execution guidance and command-completion handling. The patch and workspace instructions allow computational tasks to continue in the background while the agent responds to user feedback. When a background task finishes, the agent is automatically reactivated to inspect its outputs and continue the remaining analysis.

#### 4.1.1 Analytical skills and configured execution

Analytical methods are organized into skills covering 16S amplicon processing, reference-based metagenomic profiling, assembly-based genome reconstruction, and downstream statistical analysis, annotation and vi- sualization (table 1). Each skill contains a SKILL.md document describing its purpose, required inputs, software dependencies, reference resources, analytical procedures and expected outputs. Some skills also include reference scripts that can be adapted to the study data. These skills are registered through the plugin manifest, which defines their names, locations and metadata, and are made available through OpenClaw’s skill mechanism. For each analytical step, MicroBioAgent selects a relevant skill and reads its SKILL.md through the file-reading tool, bringing the method-specific guidance into the model’s context. This stepwise reading is guided by AGENTS.md, which specifies that the relevant skill should be consulted before execution. Software environments and reference database locations are recorded in two configuration files. The runtimes.yaml file identifies the Conda environment containing each tool or library, whereas references.yaml specifies the locations of the reference databases. The reference configuration is generated from user-supplied settings when the OpenClaw gateway starts, with database paths adjusted for its running environment. Following the resource-access guidance in TOOLS.md, the agent would read these configurations to select the required software environment and locate the reference databases. Together, modular skills and separate resource configurations allow users to extend the available analytical methods. A new method can be introduced through an additional skill that uses existing software environments and reference databases or specifies additional resources in the corresponding configuration files.

#### 4.1.2 Analysis planning and execution

MicroBioAgent uses mgx_create_job to create or reuse an analysis directory for each conversation. The tool records the association between the conversation identifier and the directory on disk, allowing successive steps to share the same locations for inputs, outputs, scripts and logs. With the analysis directory established, MicroBioAgent develops a plan based on the research objective, input data and requested outputs. The plan defines the analytical steps, their order and completion criteria, and is maintained within the conversation.

Method-specific skills provide methodological guidance for the planned analyses, while CODING.md guides script development and command execution through exec. The execution guidance standardizes agent script authoring rules to ensure operational errors are promptly captured and handled, while enabling the agent to execute independent tasks in parallel and balance concurrency and thread counts against available processor and memory resources. It also specifies how execution status, logs and generated files are checked after a command launched through exec finishes. These checks determine whether execution succeeded and whether the expected output files were produced. If either condition is not met, the agent reviews the logs and outputs to identify the problem and determine which parts of the analysis need to be rerun.

#### 4.1.3 Human-in-the-loop interaction and automatic continuation

MicroBioAgent allows users to interact with the agent while computational tasks run asynchronously in the background. Guided by CODING.md and tool instructions, the agent executes commands via exec with a 30-second initial wait period and no overall execution time limit. Commands exceeding this threshold continue running in the background, prompting the agent to invoke sessions_yield to yield its turn. MicroBioAgent handles these background processes and logs their outputs without requiring the agent to poll for status updates. This decoupling keeps the conversation responsive, enabling users to discuss or revise pending analyses without interrupting running computations. Both user messages and command-completion notifications can trigger the agent within the same thread, supporting seamless continuous interaction and automatic task continuation upon completion.

To enable automatic continuation after background execution, we developed a patch for OpenClaw (v2026.7.2) that extends its event handling and conversation activation mechanisms. Each background command is associated with its originating conversation identifier, ensuring completion events are dispatched to the corresponding conversation. For an idle conversation, the patch immediately triggers a new agent loop upon command completion without waiting for scheduled polling or user input. If an event arrives during an active agent run, it is queued and schedules a subsequent activation. This queuing mechanism prevents command results from being dropped when the agent is busy, ensuring they are reliably processed in the next turn. The injected event message contains the command result along with guidance instructions to prompt further analysis. Together with updated tool instructions that guide the agent to yield its turn, these changes allow background computation to run alongside active conversation and support automatic continuation upon task completion.

### 4.2 Case study design

To assess MicroBioAgent across diverse microbiome workflows, we evaluated the system on three published datasets spanning plant, human, and environmental ecosystems (Supplementary Table S1), covering 16S rRNA amplicon profiling, reference-based shotgun metagenomics, and assembly-based genome reconstruction. Across all three cases, the agent was powered by deepseek-v4-pro and provided with sequencing reads, sample metadata, and natural-language research goals without pre-written analysis scripts. The evaluation tested whether the agent could autonomously compose end-to-end workflows and recapitulate the primary biological conclusions of the original studies using its configured software environments and reference databases. When specific metrics or domain-standard tools were explicitly requested in the prompt, the agent applied them; otherwise, all analytical procedures, software environments, and command parameters were determined autonomously using its available skills. The specific workflows and evaluation tasks for each case study are detailed below.

#### 4.2.1 Case 1 — Tomato rhizosphere 16S and metagenomic analysis

This case evaluated MicroBioAgent’s ability to analyze paired amplicon and shotgun metagenomic datasets to identify microbial signatures associated with disease resistance in tomato. Sequencing data were retrieved from BioProject PRJNA777724 [12]. The dataset comprises 60 rhizosphere 16S rRNA (V3–V4) amplicon libraries generated from four cultivars—two susceptible (A57, HG70) and two resistant (HG64, HF12). Samples were collected across five experimental conditions: baseline (pre-inoculation at day 0), as well as inoculated and uninoculated controls at days 5 and 10 (*n* = 3 biological replicates per group). Additionally, 36 matched shotgun metagenomic libraries were profiled for the baseline (day 0) and day-5 timepoints. The evaluation comprised two analytical stages, including an initial autonomous workflow and a researcher-guided re-analysis. In the initial stage, the agent independently performed sequence processing, diversity profiling, community composition comparisons, and identification of taxa enriched in resistant cultivars relative to susceptible controls. Comparisons with the source study focused on inoculation-induced diversity shifts and the recovery of published resistance biomarkers, which were originally identified via LEfSe using an alpha cutoff of *P <* 0.1 and a logarithmic LDA score threshold *>* 2.0 [12]. In the second stage, to evaluate how profiler versions influence taxonomic recovery, the investigator prompted a follow-up reanalysis using MetaPhlAn2 to align directly with the original study’s profiling environment.

#### 4.2.2 Case 2 — Colorectal cancer faecal metagenomic analysis

This case evaluated MicroBioAgent’s ability to identify disease-associated microbial features and assess their predictive value for CRC. The analysis used the Hong Kong C1 cohort, available under accession PRJEB10878, comprising 128 faecal metagenomes from 74 patients with CRC and 54 controls [20]. The CRC group included 36 early-stage cases (stages I–II) and 38 late-stage cases (stages III–IV). The analysis tasks covered community diversity and composition, differential abundance of species and functional features, and associations with clinical variables. Functional analyses included both gene-family and KO abundance profiles. The task further required species co-occurrence network analysis to examine relationships among CRC-associated taxa, together with marker selection and classifier development to distinguish CRC cases from controls. Comparisons with the original study focused on evaluating the direction of species associations with CRC and co-occurrence patterns among disease-associated taxa, while the predictive performance of the selected markers and classifier was assessed using AUROC, sensitivity, and specificity.

#### 4.2.3 Case 3 — Polyethylene-enrichment genome-resolved metagenomic analysis

This case evaluated MicroBioAgent’s ability to recover microbial genomes from metagenomic data and characterize their functional potential. The evaluation utilized six Oxford Nanopore long-read metagenomes from project PRJEB97175, derived from two soil inocula enriched on LMWPE, triacontane, or tetracontane [23]. The analysis tasks covered metagenomic assembly, binning, quality filtration for high- and medium- quality MAGs, dereplication, and taxonomic classification, alongside cross-culture abundance profiling and gene functional annotation for MAGs assigned to ten focal taxa from the original study. Evaluation examined whether MAGs corresponding to the ten taxa of interest in the original study were recovered and whether their abundance patterns across enrichment cultures were characterized. Functional evaluation examined whether the annotations identified genes associated with the three metabolic processes investigated in the original study, namely alkane terminal oxidation, metabolism of 2-ketones or oxidized derivatives, and *β*-oxidation.

### 4.3 Robustness and ablation experiments

To reduce computational overhead during repeated testing, robustness and ablation experiments were conducted using preprocessed data from Case 1. The 16S dataset comprised the ASV table, representative sequences, a rooted phylogenetic tree, denoising statistics, and sample metadata. For shotgun sequencing, 36 quality-controlled and host-cleared libraries were used, each subsampled to 20% of its original read depth. These preprocessed inputs were used across all experimental conditions, with each session running the analyses independently.

#### 4.3.1 Robustness across models and prompt granularity

Robustness was evaluated across three large language models (glm5.3-flash, deepseek-v4-flash, and deepseek-v4-pro) and three levels of prompt detail. Each model–prompt combination was executed in three independent replicates, yielding a total of 27 experimental sessions. All three prompts requested joint 16S and shotgun metagenomic analyses of the Case 1 study, differing in how explicitly they specified the analytical steps and methods:

- **Detailed prompt:** Specified exact analytical tasks and diversity methods, covering dataset char- acterization, 16S alpha diversity (Shannon index after sequencing-depth normalization), 16S beta diversity using weighted and unweighted UniFrac and Bray–Curtis distances, weighted-UniFrac PCoA, PERMANOVA, and permutational analysis of multivariate dispersions (PERMDISP); 16S taxonomic and differential-abundance profiling, and shotgun species-level taxonomic and differential-abundance profiling. For both datasets, it requested comparing each resistant cultivar against the susceptible group at each sampling timepoint and identifying taxa enriched in both HG64 and HF12 within each of T1, T2C, and T2.
- **Medium prompt:** Retained the same analytical scope (dataset characterization, 16S diversity, 16S and shotgun taxonomic profiling, cultivar comparisons at each timepoint, and shared enriched taxa detection), but omitted specific diversity metrics and statistical tests, leaving method selection to the agent.
- **Simple prompt:** Stated only the high-level biological objective of identifying microbiome patterns associated with bacterial-wilt resistance across the 16S and shotgun datasets, without prescribing specific analytical steps or methods.

System configurations, analytical software, reference databases, and preprocessed inputs were held constant across all runs. Each replicate was initialized in a clean conversation session without further methodological guidance following initial plan confirmation.

To evaluate system performance across these conditions, task completion was systematically evaluated across six distinct analytical tasks: (1) 16S alpha diversity, (2) 16S beta diversity, (3) 16S genus-level taxonomic profiling, (4) 16S differential abundance, (5) shotgun species-level taxonomic profiling, and (6) shotgun differential abundance. A task was scored as completed when the session produced the corresponding quantitative output, such as a per-sample Shannon diversity table, a between-sample distance matrix, a genus- or species-level taxonomic abundance table, or a differential-abundance result table. For each model–prompt combination, the task completion rate was calculated as the proportion of completed runs out of the three independent sessions.

To evaluate output stability across models and prompt levels, quantitative results were averaged across completed replicate sessions within each model–prompt combination and compared across entries shared by the compared groups using two-sided Mann–Whitney *U* tests with Holm correction. For diversity analyses, per-sample Shannon indices (*n* = 60 samples) and pairwise weighted UniFrac distances (*n* = 1,770 unordered sample pairs) were averaged across replicates. For taxonomic profiling, abundance tables reported as percentages or raw counts were converted to per-sample relative abundances before removing unclassified or unnamed entries, with taxa absent from a replicate that included the sample entered as zero. Because microbial relative-abundance tables are highly right-skewed and contain many near-zero entries near the detection limit, minor numerical fluctuations in the low-abundance tail can disproportionately affect rank-based tests despite negligible differences in absolute abundance. To prevent these near-zero entries from dominating the comparison of taxonomic abundance tables, relative-abundance distributions were compared across shared sample and taxon entries with a session-mean relative abundance of at least 5% in every compared group, whereas all shared taxa (including low-abundance taxa) were included without an abundance threshold in the differential-abundance comparisons below.

For differential abundance, because the prompts did not prescribe a specific statistical test, sessions selected different methods (including DESeq2, MaAsLin 3, Wilcoxon rank-sum test, Mann–Whitney *U* test and LEfSe). Because different methods report differently defined effect sizes, log_2_ fold-change distributions were compared only across sessions that used the same statistical test. Cross-model comparisons within each prompt level used sessions that ran DESeq2 (Supplementary Table S4), and within-model comparisons across prompt levels were conducted between the detailed and medium prompts, which specified the same comparisons between resistant and susceptible cultivars at each timepoint, using the statistical test shared by both prompts for each model (Supplementary Table S5). In both comparisons, each data point represented the log_2_ fold change of a taxon for a given resistant cultivar (HG64 or HF12) and sampling timepoint relative to the susceptible control (averaging across susceptible controls within a session when A57 and HG70 were reported separately). Values were averaged across replicate sessions without imputing missing entries as zero, and entries shared across all compared groups were retained.

#### 4.3.2 Ablation of asynchronous execution and human guidance

To evaluate the role of the runtime patch for asynchronous execution and automatic continuation (section 4.1.3), we compared the complete system (full) with an ablated configuration without the runtime patch (no-patch) using deepseek-v4-pro, the detailed Case 1 prompt, and the preprocessed Case 1 dataset across three independent sessions per condition. All other system components, including microbiome domain skills, software environments, and reference databases, were held constant across both conditions.

In the end-to-end execution experiment, sessions were run with no user input after initial plan confirmation. Task completion was scored across nine required outputs (16S alpha diversity, 16S beta diversity, 16S genus- level taxonomic table, 16S differential abundance, shotgun species-level taxonomic table, shotgun differential abundance, figures, analysis scripts, and an analysis report), and method adherence was scored against the eight diversity and statistical requirements specified in the detailed prompt (sequencing-depth normalization, Shannon diversity, weighted UniFrac, unweighted UniFrac, Bray–Curtis dissimilarity, weighted-UniFrac PCoA, PERMANOVA, and PERMDISP). For each session, model call counts, tool invocations (exec, process, and sessions_yield), and total token consumption (input and output tokens, including reasoning tokens) were recorded from the session transcript.

In the asynchronous human-guidance experiment, we tested mid-run plan revision while the Case 1 QIIME 2 taxonomy-classification command was running in the background. Once the classification process was confirmed active, the user sent a standardized instruction to keep the running classification command and update the pending plan to use DESeq2 for the subsequent differential-abundance analysis and generate a heatmap of significant genera, with no further user input thereafter. Each session was evaluated on two outcomes: (1) whether the agent processed the user’s message and updated the pending analysis plan while the classification command was still running; and (2) whether, once the background classification command finished, the system automatically reactivated the waiting session so that the agent inspected the classification output and continued with the revised plan, without requiring additional user messages or repeated process calls to check whether the command had completed.

## Data availability

Raw sequencing data are accessible via BioProject accession numbers PRJNA777724, PRJEB10878, and PRJEB97175. Case-study statistics are detailed in Supplementary Data 1.

## Code availability

MicroBioAgent source code, domain procedures, agent instructions, analysis prompts, environment specifica- tions, and the session-runtime extension (distributed as a version-pinned OpenClaw patch pack) are publicly available at https://github.com/Zeming-LI-Andy/openclaw-microbiome.

## Supporting information

Results for case studies

## Supplementary Information

Case-study accessions and statistics are provided in Supplementary Data 1.

**Figure S1:**
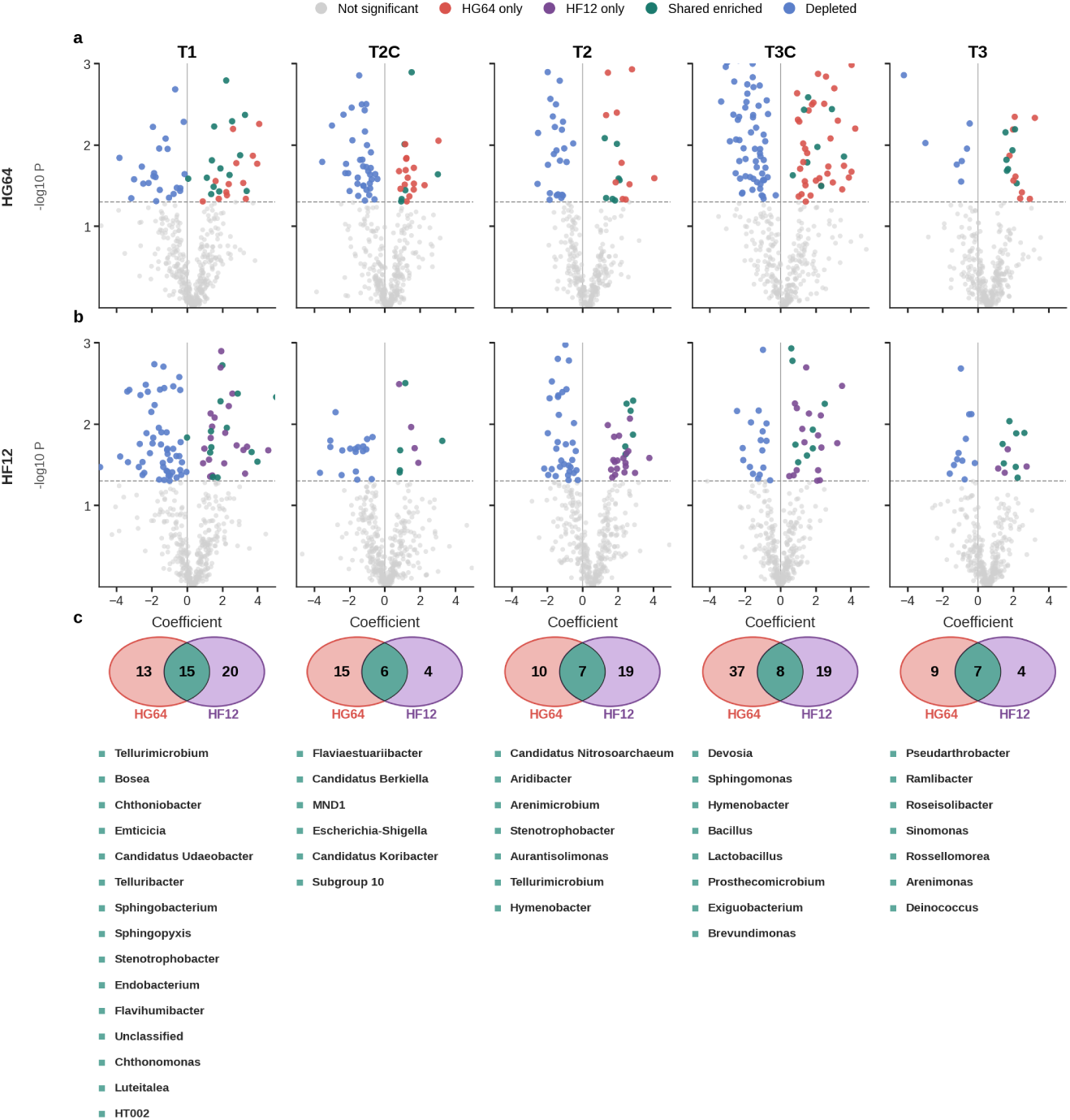
Genus-level differential abundance in the 16S profiles of the resistant tomato parent HG64 and the resistant F1 hybrid HF12. (a, b) Volcano plots of MaAsLin 3 coefficients for HG64 (a) and HF12 (b) versus the pooled susceptible cultivars at each sampling condition. (c) Overlap of genera with a positive coefficient and *P <* 0.05 in both resistant cultivars.

**Figure S2:**
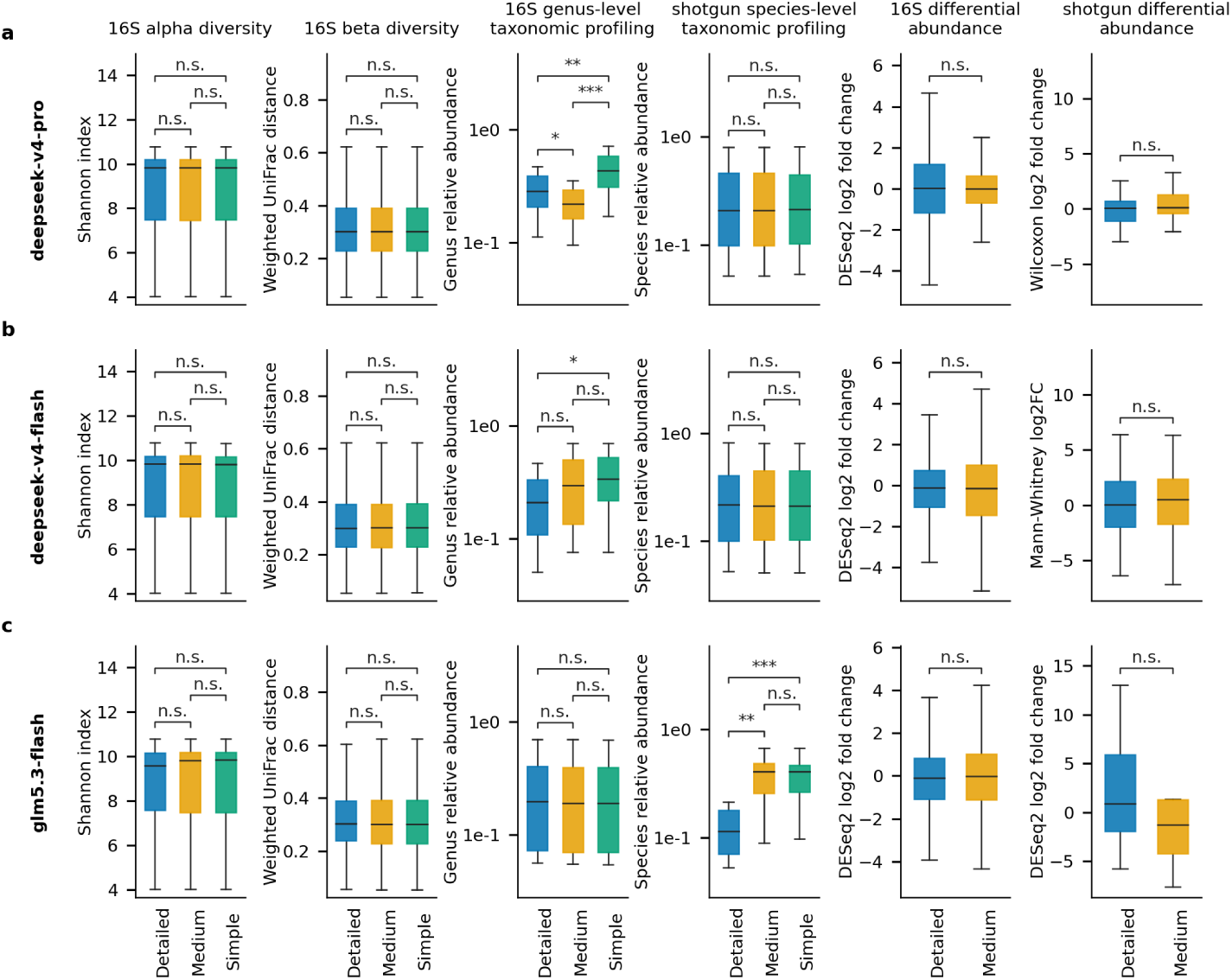
Comparison of analytical outputs across prompt levels within each model. Rows represent individual models (deepseek-v4-pro, deepseek-v4-flash, and glm5.3-flash). The first four columns display 16S Shannon diversity indices, pairwise weighted UniFrac distances, 16S genus-level relative abundances, and shotgun species-level relative abundances across prompt levels. The remaining two columns compare differential-abundance log_2_ fold changes between detailed and medium prompts only (16S: DESeq2; shotgun: model-specific tests including Wilcoxon, Mann–Whitney *U* , and DESeq2, as summarized in Supplementary Table S5). Brackets in the first four columns indicate two-sided Mann–Whitney *U* tests with Holm correction across all three prompt pairs within each model; brackets in differential-abundance panels indicate pairwise comparisons between detailed and medium prompts. \**P <* 0.05; \*\**P <* 0.01; \*\*\**P <* 0.001; n.s., not significant (adjusted *P ≥* 0.05). Values are derived from repeated analyses of the same Case 1 dataset and do not represent independent biological replicates.

**Table S1:** Overview of the three case-study datasets.

| Case | BioProject | Sequencing libraries | Workflow |
|---|---|---|---|
| Tomato rhizosphere | PRJNA777724 | 60 16S; 36 matched shotgun | 16S and shotgun profiling |
| CRC faecal metagenomes | PRJEB10878 | 128 shotgun | Taxonomic and functional profiling |
| Polyethylene enrichment | PRJEB97175 | 6 long-read shotgun | Genome-resolved metagenomics |

**Table S2:** Principal software tools and reference databases used across the three case studies.

| Software or reference database | Version or release |
|---|---|
| QIIME 2 | 2024.10.1 |
| DADA2 | 1.30.0; QIIME 2 plugin 2024.10.0 |
| SILVA 138.2 | 138.2 |
| KneadData | 0.12.4; hg37dec_v0.1 |
| MetaPhlAn | 3.0.7; mpa_v30_CHOCOPhlAn_201901 |
| MetaPhlAn | 2; mpa_v20_m200 |
| HUMAnN | 3.0.0.alpha.4; ChocoPhlAn v296_201901 |
| MaAsLin3 | 1.2.0 |
| Aviary | 0.13.2; assembler myloasm 0.3.0 |
| CheckM2 | 1.1.0 |
| CoverM | 0.7.0 |
| GTDB-Tk | 2.7.2; database R232 |
| eggNOG-mapper | 2.1.13; eggNOG 5.0.2 |

**Table S3:** Comparison of reported capabilities across bioinformatics agents for end-to-end microbiome analysis from raw sequencing reads.

|  | Agent operation |  | Microbiome analysis from reads |  |  | Deployment |
|---|---|---|---|---|---|---|
| Agent | Autonomous workflow execution | Human-in-the-loop | 16S amplicon | Reference-based shotgun | Assembly / MAG | On-premises analysis |
| BIA | ✓ | × | × | × | × | ✓ |
| BioChatter | × | ✓ | × | × | × | ✓ |
| AutoBA | ✓ | × | × | × | × | ✓ |
| Biomni (open-source) | ✓ | × | × | × | × | ✓ |
| BioAgents | × | × | × | × | × | × |
| M3A | ✓ | ✓ | × | × | × | ✓ |
| BioMedAgent | ✓ | × | × | × | × | ✓ |
| BioMaster | ✓ | × | × | ✓ | × | ✓ |
| MetaClaw | ✓ | × | ✓ | ✓ | ✓ | ✓* |
| MicroBioAgent | ✓ | ✓ | ✓ | ✓ | ✓ | ✓ |

*Notes:* This comparison evaluates the functional scope of existing bioinformatics agents based on published literature and open-source repositories, highlighting MicroBioAgent’s ability to unify autonomous raw-read execution, interactive human guidance, and local execution across all major microbiome workflows. ✓, supported; ×, not supported. \*Upstream processing requires cloud-based FlowHub, while downstream tasks run locally. MAG, metagenome-assembled genome.

**Table S4:** Replicate sessions and shared entries for cross-model DESeq2 comparisons.

| Dataset | Prompt | DeepSeek Pro | DeepSeek Flash | GLM Flash | Shared entries |
|---|---|---|---|---|---|
| 16S rRNA gene, genus | Detailed | 1/3 | 2/3 | 3/3 | 1,724 |
| 16S rRNA gene, genus | Medium | 1/3 | 2/3 | 1/3 | 2,659 |
| Shotgun metagenome, species | Detailed | 1/3 | 2/3 | 2/3 | 228 |

*Notes:* Values for each model indicate the number of independent replicates (out of three) that selected DESeq2 for differential-abundance testing. Shared entries denote the number of overlapping taxa evaluated across all three models. Shotgun metagenome results under the medium prompt were not compared because no single differential-abundance method was selected by all three models.

**Table S5:** Replicate sessions and shared entries for prompt-level differential-abundance comparisons.

| Model | Method | Detailed sessions | Medium sessions | Shared entries |
|---|---|---|---|---|
| *16S rRNA gene, genus* |  |  |  |  |
| deepseek-v4-pro | DESeq2 | 1/3 | 1/3 | 891 |
| deepseek-v4-flash | DESeq2 | 2/3 | 2/3 | 5,108 |
| glm5.3-flash | DESeq2 | 3/3 | 1/3 | 3,984 |
| *Shotgun metagenome, species* |  |  |  |  |
| deepseek-v4-pro | Wilcoxon | 1/3 | 1/3 | 39 |
| deepseek-v4-flash | Mann–Whitney *U* | 1/3 | 1/3 | 120 |
| glm5.3-flash | DESeq2 | 2/3 | 1/3 | 52 |

*Notes:* Session counts indicate the number of independent replicates (out of three) that selected the specified statistical method under each prompt level. Shared entries denote the number of overlapping taxa evaluated across both prompt levels.

**Table S6:** Ablation study of the runtime patch on end-to-end execution efficiency and token consumption.

| Condition | Session | Outputs (of 9) | Methods (of 8) | Model calls | exec calls | process polls | yield calls | Total tokens (×10<sup>6</sup>) |
|---|---|---|---|---|---|---|---|---|
| full | Replicate 1 | 9/9 | 8/8 | 150 | 91 | 1 | 17 | 12.92 |
| full | Replicate 2 | 9/9 | 8/8 | 124 | 74 | 21 | 5 | 14.81 |
| full | Replicate 3 | 9/9 | 8/8 | 126 | 82 | 0 | 13 | 15.84 |
| full | Mean / sum | 27/27 | 24/24 | 133.3 | 82.3 | 7.3 | 11.7 | 14.53 |
| no-patch | Replicate 1 | 8/9 | 8/8 | 211 | 77 | 102 | 0 | 23.77 |
| no-patch | Replicate 2 | 9/9 | 8/8 | 403 | 142 | 227 | 0 | 49.32 |
| no-patch | Replicate 3 | 9/9 | 8/8 | 245 | 99 | 136 | 0 | 32.15 |
| no-patch | Mean / sum | 26/27 | 24/24 | 286.3 | 106.0 | 155.0 | 0.0 | 35.08 |

*Notes:* full and no-patch denote the system with and without the runtime patch, respectively. All runs were conducted using deepseek-v4-pro with the detailed Case 1 prompt on the preprocessed Case 1 dataset. Summary rows report pooled counts for outputs and methods across the three replicates, whereas tool-call counts and token consumption are reported as per-session means. Total tokens represent the sum of input and output tokens (including reasoning tokens).

**Table S7:** Ablation study of the runtime patch on asynchronous human guidance and automatic session reactivation.

| Condition | Plan updated while<br>command running | Automatic reactivation<br>upon command completion |
|---|---|---|
| full | 3/3 | 3/3 |
| no-patch | 0/3 | 0/3 |

*Notes:* full and no-patch denote the system with and without the runtime patch, respectively. ‘Plan updated while command running’ indicates whether the agent successfully incorporated mid-run user instructions into the pending analysis plan during background execution. ‘Automatic reactivation upon command completion’ indicates whether the waiting session automatically resumed upon background task completion to proceed with the revised plan, without requiring additional user prompts. Results are reported as successful sessions out of three (*n* = 3).

